# Nematic order in cellular tissues: a standardized framework and anomalous defect dynamics

**DOI:** 10.64898/2026.04.22.719598

**Authors:** Nicolas Rembert, Mathieu Dedenon, Aurélien Roux, Claire A. Dessalles

## Abstract

Cellular monolayers often exhibit orientational order, with nematic alignment of cell shape and cytoskeletal structures governing tissue-scale collective dynamics. Despite extensive studies, a unified analysis framework for characterizing active nematics in living systems remains partial, and key discrepancies with theory persist. Here, we present a systematic and comparative analysis of nematic order and tissue flow dynamics across twelve distinct cell types. We quantify the impact of analysis parameters and provide data-driven guidelines to improve reproducibility and cross-study comparability. Across all nematic systems, we uncover remarkably consistent static properties, supporting the universality of nematic behavior in living tissues. By combining orientation-field analysis with velocity-field measurements and numerical simulations, we show that all examined systems display contractile active nematic signatures, with characteristic flow structures around topological defects. However, direct tracking of individual defects reveals subdiffusive dynamics, in stark contrast with the superdiffusive, self-propelled motion predicted by the hydrodynamic theory of active nematics. Our results establish a standardized framework for nematic analysis in biological systems and highlight fundamental limitations of current active nematic models in describing defect dynamics in living tissues.

## I. INTRODUCTION

The liquid crystal phase is a state of matter where particles display orientational order without positional order [1]. Cellular assemblies can behave as liquid crystals, exhibiting long range orientational order [2–10]. This order can be found in the orientation field of the anisotropic particles, such as the cell or cytoskeleton filaments [11, 12], where order is typically nematic. It can also be found in the velocity field of collectively migrating cells [13–15], where order is mostly polar. Liquid crystal phases have been found in cultured cellular assemblies as well as in developing tissues, where they orchestrate multiple biological phenomena, from single cell events, such as apoptosis or differentiation, to whole organism morphogenesis [16–20]. Topological defects - points or regions where the order is lost - play a key role in these processes through their role in organizing active stresses. Although numerous studies have now investigated the nematic nature of cellular tissues, a unified and consolidated view is lacking. This difficulty can be attributed to various factors, among which we identified two critical points: one, the absence of standard image analysis procedures and quality criteria to extract the orientation fields and identify topological defects; two, reports of experimental observations that challenge the theoretical interpretations within the active nematics framework [21–23].

For the first point, the main hurdle lies in the discrete nature of experimental data when compared to continuous theory. Indeed, numerous user-defined parameters are required during a nematic analysis pipeline. Their impact on the quantifications has not been thoroughly investigated, and their values are seldom reported in the literature. As a result, the interpretation of the computed nematic parameters or comparisons between studies are difficult.

For the second point, the more data is published the more evident it becomes that the framework of active nematics needs to be expanded to better characterize living cell tissues. The theory of active nematics [24– 26] successfully predicted spontaneous flows in confining channels and self-propulsion of +1/2-topological defects based on active nematic stresses, confirmed by experiments with in vitro cell tissues [7–9]. Nevertheless, tissues of a given cell type can be described as extensile or contractile nematics depending on experimental conditions and assumptions on theoretical parameters [7, 19, 23, 27–29]. To account for this apparent discrepancy, theory extensions recently introduced the coupling of two nematic fields [21, 22, 30], either adding non-linear active forces [23], or including compressibility and density-nematic couplings [31]. This on-going effort relies on a dataset that is distributed amongst multiple studies, as one study usually focuses on a specific aspect and metric. A self-consistent, quantitative, and complete characterization of nematic and dynamic properties of different cellular systems is still lacking.

Here, we characterized the nematics and dynamics of twelve different cell types (see supplementary Fig. S0). First, we propose a method to discriminate between isotropic and nematic phase in cellular monolayers. We then systematically investigated the contribution of each user-defined parameter in the analysis and propose a rationale for their choice. We computed various nematic metrics for all the nematic cell types and monitored topological defect dynamics, revealing signatures of sub-diffusive dynamics from weakly contractile active units. In contrast, the active nematic theory predicts super-diffusive defect dynamics in all the conditions explored, indicating a key missing ingredient. Finally, we extracted cell velocity fields and investigated their correlation to the nematic field, further showing that all the nematic cell types explored behave as contractile active fluids, with flow and divergence patterns around defects consistent with theoretical predictions.

## II. RESULTS

### A. Cellular monolayers can behave as nematic phases

To probe the universality and variability of nematic properties across cellular monolayers, we selected twelve different cell types (muscle cells, fibroblasts, epithelial cells and cancer cells) and cultured them on tissue culture treated 6 well plate. Monolayers were imaged overnight in phase contrast brightfield twice, at confluency and 48h later, after which they were fixed, stained for actin and nuclei and imaged. The cell orientation and velocities were then extracted from the movies and analysed.

To quantify the nematic nature of a cellular tissue, the first step is to extract the local orientation of the mesoscopic particle. This can be achieved by segmenting single cells or filaments using machine learning algorithms [32] and computing their individual orientation to get a tissue wide director field; or by using gradients of intensity. We will focus here on the latter, as the former has been extensively described and is not specific to nematic analysis [33]. Intensity gradients are common features of any microscopy image, such as brightfield or fluorescent images. They can be used as a proxy for particle orientation: gradients are sharper in the direction orthogonal to the particle axis. The gold standard tool for this approach is OrientationJ, a FIJI plugin or its python equivalent OrientationPy. Both require two user defined parameters: the tensor window size and the grid size, also called downsampling factor (Fig 1A).

**FIG. 0:**
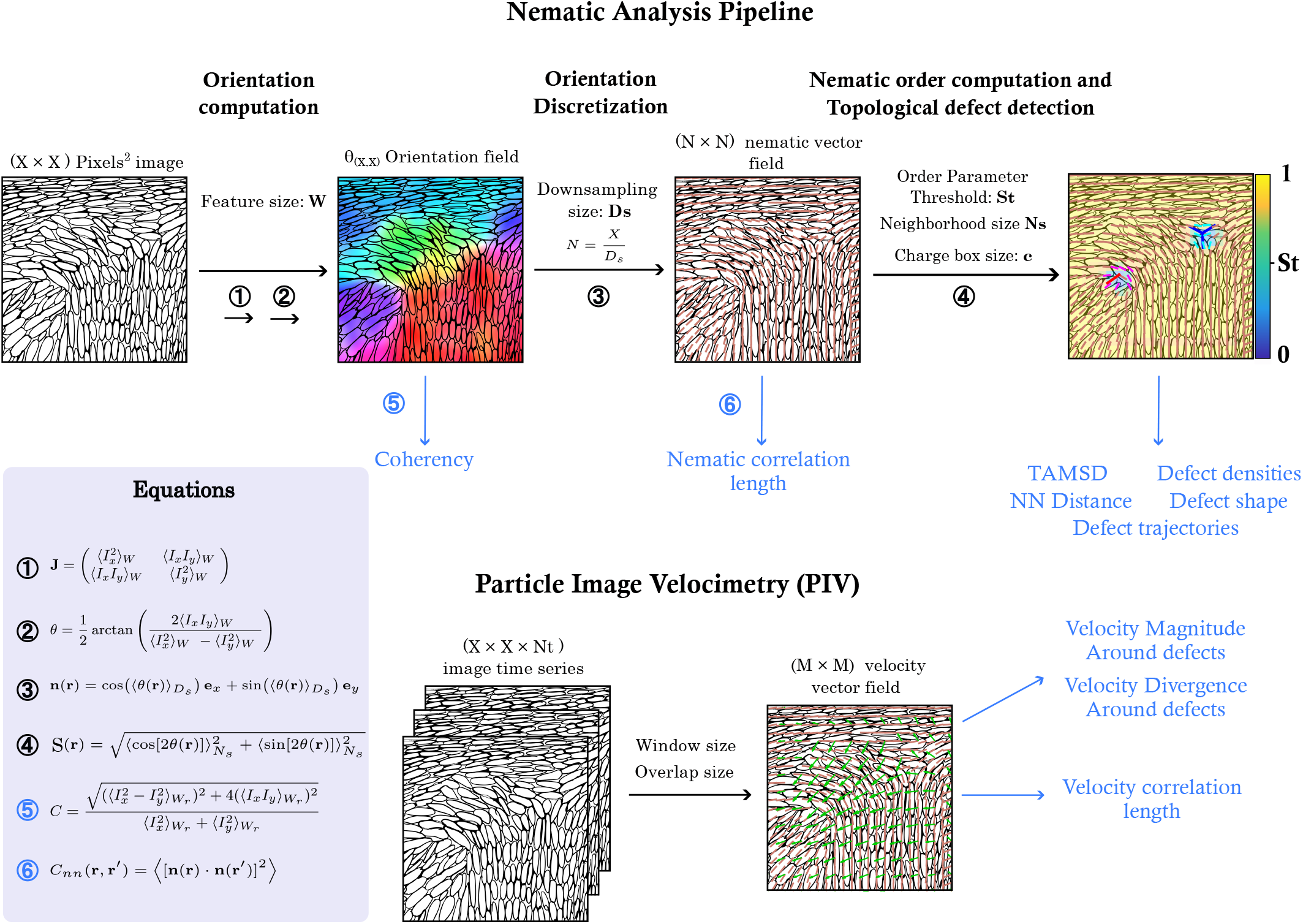
Graphical abstract: User-parameterized image-analysis pipeline extracting orientation and velocity fields from time-lapse images and producing quantitative outputs including nematic order, topological defects, and their dynamics.

**FIG. 1:**
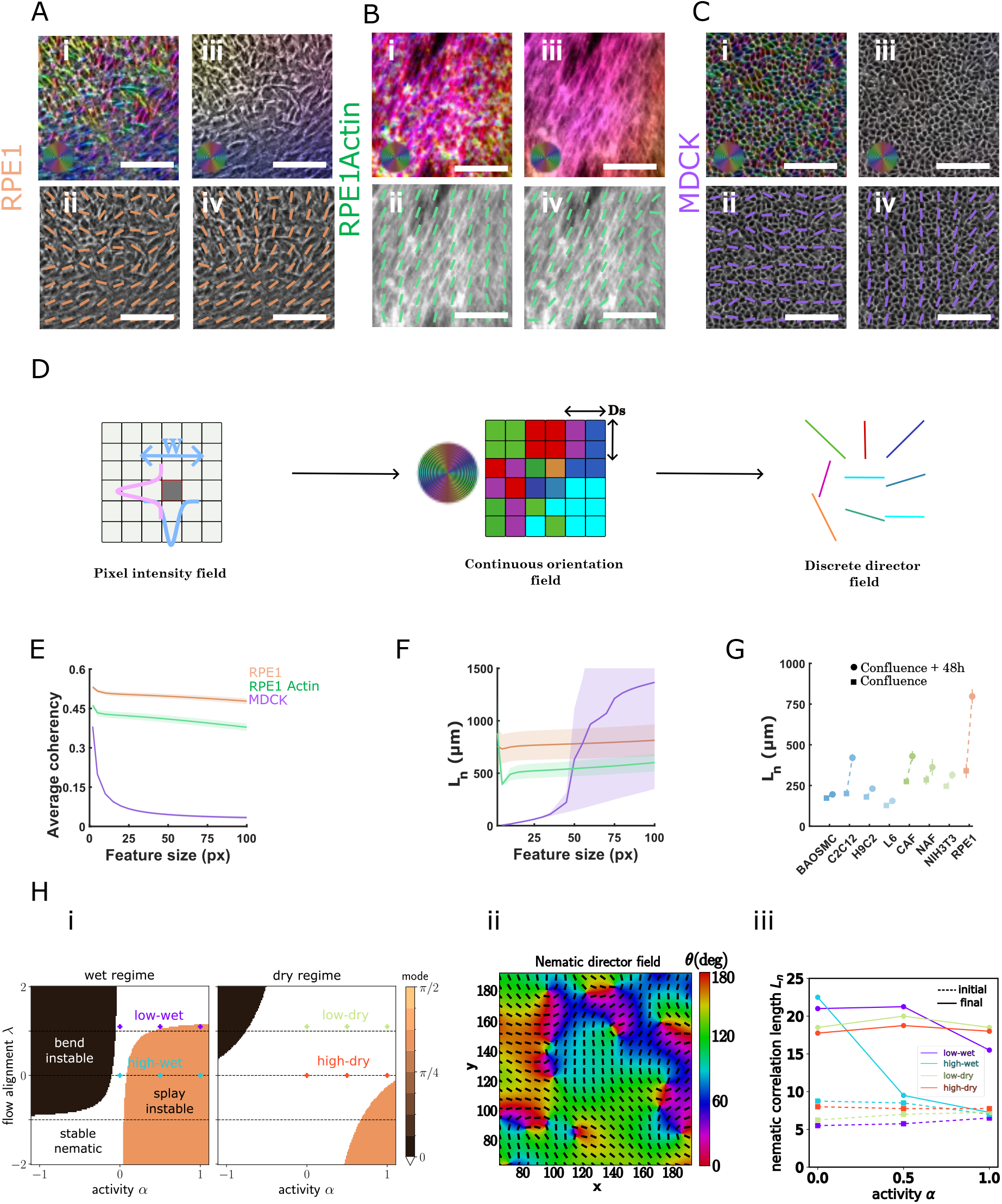
Orientation analysis of cellular monolayers after 4–5 days of culture for brightfield images of RPE1 cells (**A**), fluorescent images of RPE1 actin network (**B**) and brightfield images of MDCK cells (**C**), showing the orientation colour map (**i**,**iii**) and corresponding director field (**ii**,**iv**), for a feature size of 2 pixels (**i**,**ii**) and 50 pixels (**iii**,**iv**) and a down-sampling size of 50 pixels. Scale bar, 100 *µ*m (**A, B, C**).(**(D)**) Construction of the local orientation field from the pixel intensity field. Spatial gradients *I*_*x*_ = ∂*I/*∂*x* and *I*_*y*_ = ∂*I/*∂*y* are computed within a window *W* to build the structure tensor *J*. The local orientation *θ*(**r**) is obtained from the tensor components *J*_*ij*_, yielding a continuous orientation field. Averaging the orientation over a down-sampling size *D*_*s*_ produces the discrete director field **n**(**r**). (**E**). Average coherency as a function of the feature size used for orientation measurement. Shaded regions represent the standard deviation. Purple denotes MDCK cells, green denotes actin-based RPE1 measurements, and orange denotes RPE1 cell orientation. (**F**). Correlation length of the director field as a function of the feature size used for orientation measurement. Correlation lengths were computed using Equation 1 from N=3 images. (**G**). Correlation length measured at confluence (squares) and 2 days after confluence (circles) for all nematic cell types analyzed, feature size 50 pixels=32 *µ*m. (1 pixel=0.65*µ*m). (**H**). (**i**) Theoretical parameter space as a function of activity and flow alignment, without substrate friction (wet) and with substrate friction (dry). The regimes studied are indicated in purple (low-wet), blue (high-wet), green (low-dry) and red (high-dry). (**ii**) Snapshot of a numerical resolution of hydrodynamic equations showing the nematic director field, for the ‘high-dry’ regime at *α* = 0.5. (**iii**) Nematic correlation length as a function of activity for the different regimes. Full (dashed) lines indicate initial (final) time.

For each pixel on the image, a local orientation is calculated on the gaussian analysis window defined by the tensor window size (Fig 1 A,B,C,D). The tensor window size determines the size of the mesoscopic particle, which sets a typical lengthscale of the nematic field of interest, and thus corresponds to the feature size. Increasing the feature size is similar to a coarse-graining method, smoothing out local variations. For an isotropic tissue, increasing the feature size leads to a seemingly ordered tissue with uniform orientation, due to the averaging of random orientation (Fig 1C). To avoid the mischaracterization of isotropic tissue as nematic, we recommend assessing coherency, a measure of the strength of the local gradients (see Methods IV R). In isotropic tissues, the coherency drops to near zero when increasing the feature size, while in nematic tissues, after a short drop, the coherency remains nearly constant (Fig 1E, S1A).

The autocorrelation length is defined as the decay length obtained from an exponential fit of the nematic orientational correlation function (see Methods) and is a measure of the nematic length scale. Its evolution as a function of feature size differs between isotropic and nematic tissues: in the latter, it is near constant as there is an intrinsic nematic length scale in the tissue, while in the former, it increases artificially due to noise averaging. OrientationJ performs vector averaging instead of tensorial averaging: an isotropic vector distribution becomes a spatially uniform field oriented along 0 due to the periodicity of angles. Large autocorrelation nematic lengths can therefore be found in isotropic tissues with low coherency, but they are meaningless as there is no nematic phase.

Cellular tissues can have local fluctuations of orientation within a long-range order, for instance due to cell division that still maintain local nematicity. Depending on the biological phenomenon of interest, different window sizes can be used to capture the nematic texture at the corresponding lengthscale (Fig 1E). For each new cell type, a rapid evaluation of the variations of the coherency and autocorrelation length as a function of feature size allows the selection of a well-adjusted feature size value (Fig 1E). Here, for all further quantifications we chose a 50 pixels (32 *µ*m) feature size.

The downsampling factor corresponds to the size of the square of pixels that are averaged together to form one grid point, thereby subsampling the pixels-based orientation obtained at the previous step (Fig 1D). All further parameters are calculated on this discrete grid. As a nematic field is theoretically a continuum, the smallest grid sizes lead to a finer discretization and more accurate subsequent quantifications (Fig. S1C S2E, S3A-B). The lower bound for the downsampling factor is set by the computation time. Increasing the downsampling factor also leads to a coarse graining effect, similar to increasing the window size, but here the effect is based on discretization and averaging rather than a physical phenomenon (Fig. S2E). We therefore suggest using the window size to select for the desired nematic lengthscale and the smallest downsampling factor to reduce numerical errors induced by the discretization of a continuous field. Although the correlation length is fairly independent from the down sampling factor (Fig. S1C), other quantifications such as defect density and defect trajectories are highly sensitive to it. Based on the defect trajectory results (Fig 4), the down sampling factor is set here to 10 pixels (6.5 *µ*m). Evaluating the coherency and correlation length for all eight cell types shows a clear distinction between two groups: HBEC and MDCK epithelial cells, U2OS and SKMEL cancer cells are isotropic, while C2C12 myoblasts, H9C2 myocytes, L6 myoblasts, NAF, NIH3T3 and CAF fibroblasts and RPE1 epithelial cells are nematic (Fig. S1A). Across the nematic cells, large differences are observed in correlation length, from 200 to 800 microns, uncorrelated to cell size (Fig. S1B). On average, nematic domains span across tens of cells for all nematic cell types. A similar level of order was observed in the supra-cellular organization of actin, with correlation lengths on the same order of magnitude as those measured for cellular nematic domains (Fig. S1E–F). Within one cell type, density also affects the coherency and the autocorrelation length: 48h after confluency, the correlation length increases in all nematic tissues, in particular in C2C12 and RPE1 cells (Fig 1G), suggesting that these tissues get more ordered with time.

To analyze experimental results within an appropriate theoretical framework, we solve hydrodynamic equations of active nematics for two-dimensional velocity and nematic fields (see theoretical methods in Sec. IV.R). Periodic boundary conditions and two-dimensional incompressibility are assumed, neglecting cell proliferation (Fig 1H ii). The nematic defect core size is used as a length unit to adimensionalize equations (Sec. IV.R). The system dynamics becomes a function of four dimensionless parameters, the active stress *α*, the flow-alignment *λ*, the viscosity *η*, and the substrate friction *ξ*. These parameters define two length scales, active 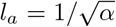 and hydrodynamic 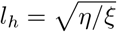. Experimental findings on Fig 5 are compatible with contractile active stress, fixing the sign *α >* 0. From linear stability analysis (Sec. IV.R), a uniform nematic order is destabilized when the product *α*(*λ* ± 1) reaches a threshold value, Fig 1Hi. For a given contractile activity, the flow-alignment parameter then defines low-activity (high-activity) regimes when the system is linearly stable (unstable). The ‘high-activity’ regime corresponds to ‘active turbulence’[34]. We also test the influence of substrate friction *ξ*, defining ‘wet’ and ‘dry’ regimes when it is absent or present, respectively. Thus, we investigate four regimes ‘low-wet’, ‘low-dry’, ‘high-wet’ and ‘high-dry’ (Fig 1Hi) with parametric values given in Sec. IV.R.

Numerical solutions are initialized with a completely disordered state, to mimic the density-dependent transition from isotropic to nematic state upon tissue confluence [10]. Then, the theoretical onset of nematic ordering is identified with experimental tissue confluence (Fig. S1D i). The numerical solution evolves until 90% of defects have been annihilate without activity, corresponding to *t*_*f*_ = 100 in dimensionless units (Fig. S1D ii). The nematic correlation length *L*_*n*_ grows with time and depends on ordering time, activity *α* and the defined regimes (Fig 1Giii). In the ‘high-wet’ regime, *L*_*n*_ converges towards the active length *l*_*a*_ at high activity without time delay. Instead for ‘low’ and ‘high-dry’ regimes, *L*_*n*_ is less sensitive to activity and exhibits a significant ordering from initial to final time. Based on experimental measurements of nematic correlation length (Fig 1F), one expects that most cell types investigated here are not in a high-activity regime, far from any turbulent phase.

### B. Half integer topological defects are found in nematic cellular tissues

The order parameter quantifies the intensity of the order within the tissue and allows to isolate defects. It is computed from the orientational field, obtained from either the segmentation or gradient of intensity method, using the equations in Methods IV G (Nematic order computation). The order parameter is calculated on the neighbourhood of a finite size around each sampling point, due to the discrete nature of the nematic analysis. The neighbourhood size is the third user-defined parameter. It should be as small as possible to match the continuum theory. Here it was set to its minimum value, i.e. the downsampling size: 10 pixels, using only the first neighbour in each direction. Increasing the neighbourhood size leads to a reduction of the order parameter variations: ordered regions have lower order and disordered “wells” have higher order and are wider (Fig 2A).

**FIG. 2:**
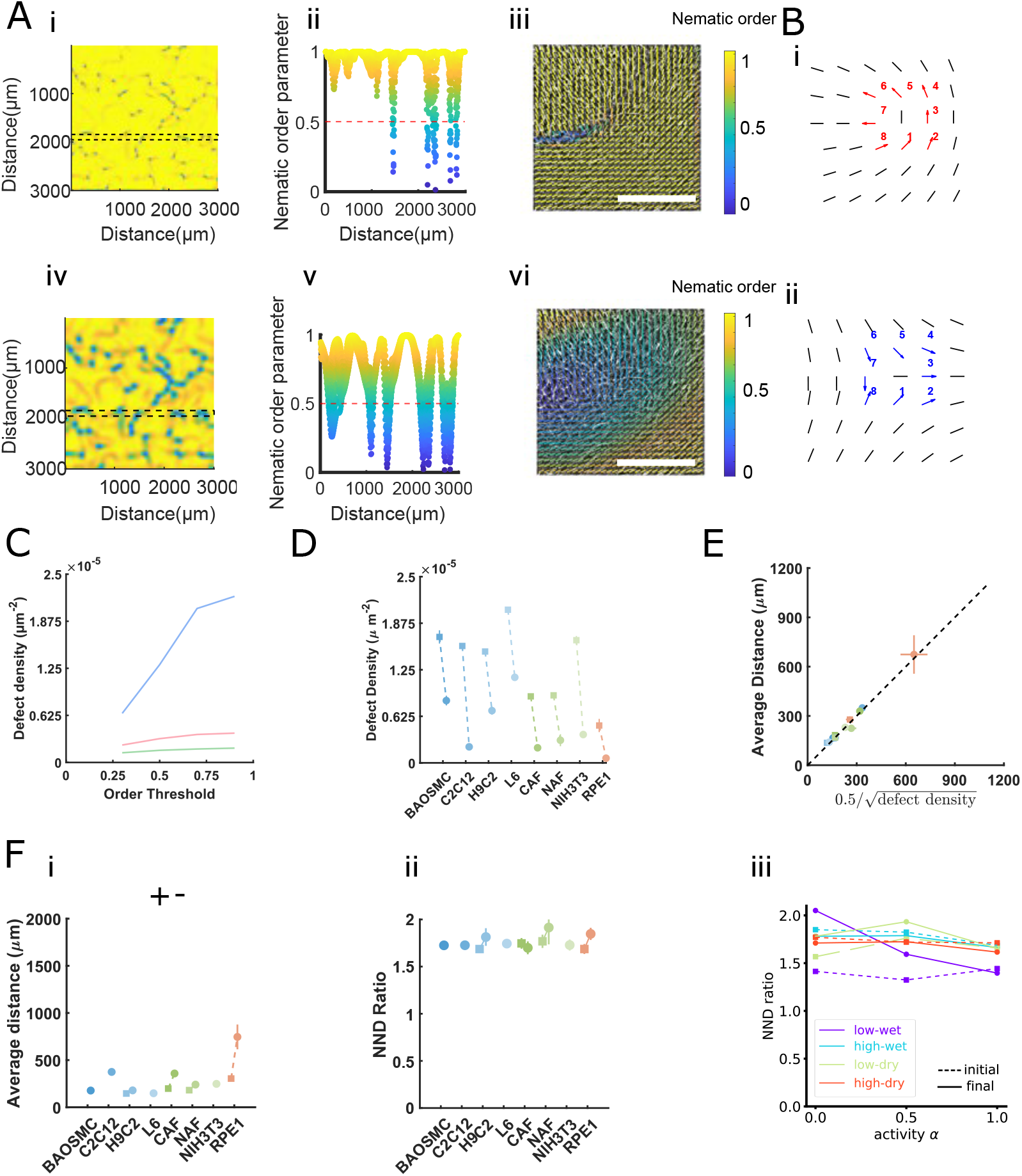
(**A**). Spatial characterization of nematic order. (**i**) Nematic order color map computed using Equation 2 with a neighboring distance of 60 pixels. Dotted lines indicate the region used to extract the profile shown in (**ii**). (**ii**) One-dimensional profile of the nematic order parameter, showing regions of low order. The dotted horizontal line at 0.5 denotes the threshold used to define low nematic order (**iii**) Director field computed with an orientation window size of 50 pixels and a grid spacing of 30 pixels, with vectors color-coded by the local nematic order parameter. Scale bars show 100 *µ*m (**iv–vi**) Nematic order color map, nematic order profile, and director field computed using Equation 6 with a neighboring distance of 180 pixels. (**B**). Schematic of the topological defect detection based on the computation of a discrete winding number. (**C**). Topological defect density (*µ*m^−2^) as a function of the nematic order threshold value (*S*_*threshold*_) for a fixed down sampling size and varying orientation-analysis feature sizes (blue 5 pixels, red 20 pixels, green 100 pixels). (**D**). Topological defect density measured at confluence (squares) and 2 days after confluence (circles) for all nematic cell types analyzed. (**E**). Average distance of nearest neighbor defects as a function of 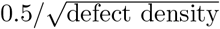. (**F**). Average nearest-neighbor distance (NND) between defects of opposite topological charge (**i**). Ratio between the NND of same-charge to opposite-charge defects, for experiments (**ii**) and theory (**iii**).

Theoretical topological defects are defined as points where the order parameter is zero. In discrete analysis, they are selected from points where the order parameter is below a certain user-defined threshold, set to 0.5 here, defining a fourth parameter (Fig 2A ii, v). Increasing the order parameter threshold leads to more defects as wells of higher order are included (Fig 2C). These points are further filtered by their topological charge, computed as the winding number. The winding number is the integral of the nematic vector variations along a circle of a userdefined distance. This distance is called the charge box size, set to one grid size here, defining a fifth parameter (Fig 2B). To calculate the winding number, only defects with neighbours further away than the charge box size were used to avoids errors due to a close defect that would perturb the nematic field in the vicinity of the defect of interest. Points where the winding number is equal to +1/2 or −1/2 are kept and defined as defects. Because the low order patches span multiple points, defects are further filtered to keep only one defect per well of low order, by selecting the center of mass of the connected defects of same charge (see Methods).

The density of detected defects depends on the feature size and on the downsampling factor (Fig 2C, S2F). As smaller feature sizes capture local fluctuations in the nematic order, the number of defects increases (Fig 2.C, S2B).In isotropic tissues, such as MDCK monolayers, multiple large regions of low order can naturally appear. This effect is further accentuated when the feature size is small, as the orientation computation captures small orientation variations within the tissue, particularly at cell membranes. As a result, extended disordered wells may emerge and lead to an artificially high density of detected defects (Fig. S2A). In nematic tissues, a net separation between regions of low and high nematic order can be done, reflecting large scale orientational properties similarly to the correlation length (Fig 1E). As the feature size is increased, nearby defects merge or annihilate to keep only a large-scale nematic texture characterizing the tissue organization.

The density of defects differs between cell types, following the inverse variations as the autocorrelation length as expected, with for instance fewer defects in C2C12 and RPE1 (Fig 2D). It did not scale with the cell size (or aspect ratio), showing that nematic properties are linked to specific cell properties and not simply scaling with size (Fig. S2C). The number of defects drops with time, as the tissue gets more ordered, consistent with the increase of the nematic correlation length (Fig 1, S2D). Overall, defects for all nematic cell types follow a Poisson’s distribution, with the average inter-defect distance scaling with the defect density to the power −1/2 (Fig 2E). Separating defects by their charge shows that their ratio is one, leading to a vanishing total charge, as expected for a flat infinite domain (Fig. S2E).

Furthermore, the nearest-neighbour distance among defects of the same charge is roughly twice the nearestneighbour distance to a defect of opposing charge, suggesting that the defect charges are spatially correlated and their distribution is not an independently marked Poisson process (Fig 2F). Instead, it suggests short range pairing of oppositely charged defects and repulsion between similarly charged defects, typical of a 2D Coulomb-gas of nematic defects. This is confirmed by numerical results of active nematohydrodynamics (Fig 2F), where the ratio between nearest-neighbour distances of defects remains between 5/4 and 2 with weak sensitivity to the investigated regimes. Similarly to the nematic correlation length, this ratio is weakly dependent on time at high activity but increases over time for low activity.

### C. Topological defect shape informs on mechanical properties of tissues

Individual topological defect found in nematic cellular tissues vary in shape, either more U-shaped, with more bend, or V-shaped, with more splay (Fig 3. A i ii) This could reflect different mechanical properties: resistance to bend (encoded by the elastic constant *K*_1_) and to splay (*K*_3_), which can be evaluated from the average defect shape [3, 35, 36]. By computing how the vector orientation varies when rotating around +1/2 defects and comparing this curve to the simulated ones we can extract the *K*_1_*/K*_3_ ratio. Although individual defects show shape-variations, for the six cell types, the average defect corresponds to the *K*_1_ = *K*_3_ case, as hypothesized in most simulations work (Fig 3B). Performing the same analysis on theoretical data for which *K*_1_ = *K*_3_ is imposed explicitly in the equations, we find an important variability of the +1/2 defect shapes. For high activity, the hydrodynamic flows deform +1/2 defects into splay-dominant (Fig 3Ci) or bend-dominant (Fig 3Cii) shapes. The different regimes influence defect shape in a non-trivial way, with splay-dominant shape only for the ‘high-wet’ regime (Fig 3Ciii). By computing the integrated difference of the angular profile from the linear profile expected at *K*_1_ = *K*_3_, we construct a dimensionless shape anisotropy *ε* taking extreme values for perfect splay or bend shapes. As a function of activity, shape anisotropy is strongly dependent on activity amplitude with important individual defect variability (Fig 3Civ). Therefore, a comparison to experiments indicates that the cell tissues should be in a low activity regime (Fig 3C). In general, the analysis of the shapes of the defects +1/2 is a non-invasive tool to infer the mechanical properties of the nematic tissues.

**FIG. 3:**
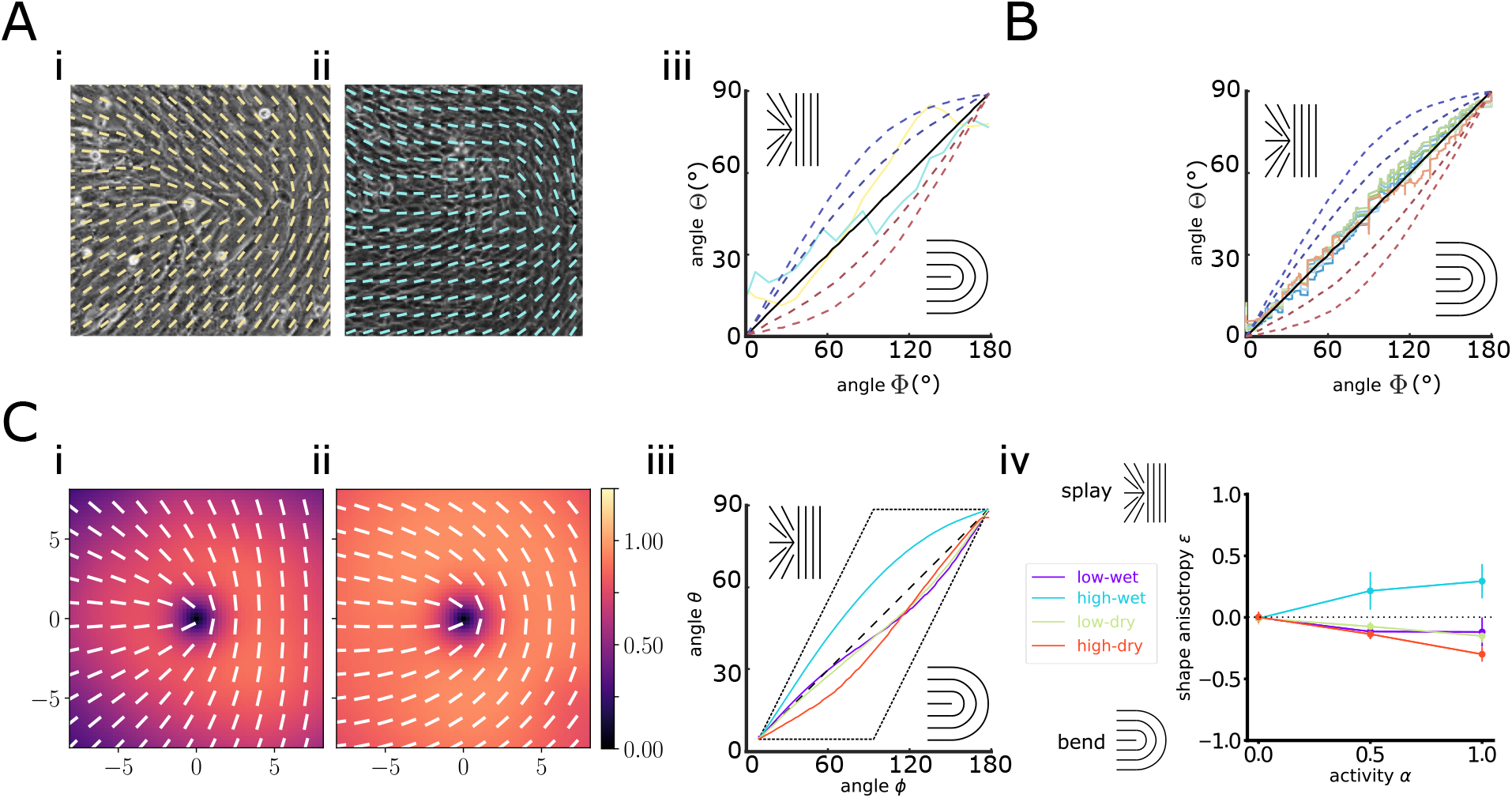
Topological defects shape arises from tissue mechanical properties (**A**). Characterization of single +1/2-topological defects shape. Scale bars show 100 *µ*m (**i**) H9C2 cell line. (**ii**) L6 cell line showing the +1/2-topological defect director field. (**iii**)Angular distribution describing the shape of these defects (see methods for computation of distribution). (**B**). Angular distribution describing the average shape of +1/2-topological defects for different cell types (color code as in (Figure 1 (**G**)). Solid curves correspond to theoretical predictions for different ratios of Frank elastic constants: *K*_1_ = *K*_3_ (black), *K*_1_ *> K*_3_ (red), and *K*_3_ *> K*_1_ (blue). (**C**). Shape of +1/2 topological defects in numerical solutions with *K*_1_ = *K*_3_. Extreme cases for splay-dominated (**i**) or bend-dominated (**ii**) shapes, corresponding respectively to high-wet and high-dry regimes at *α* = 1. The color scale indicates the nematic order field (**i**,**ii**). (**iii**) Average angular distribution for the different theoretical regimes for activity *α* = 1, with dashed lines corresponding to perfect splay (left) or perfect bend (right). (**iv**) Quantification of the shape anisotropy as a function of activity in the different regimes, where *ϵ* = +1 (*ϵ* = −1) corresponds to perfect splay (bend).

### D. Topological defects have sub-diffusive dynamics

A decreasing number of defects with time is a hallmark of passive nematic fluids, where defects are initiated during the isotropic-nematic transition, attract each other and annihilate progressively. In contrast, in turbulent active nematics, defects are actively created and annihilated with time, keeping a constant density at steadystate [34]. We therefore investigated defect dynamics in these cellular systems where time also correlates with increasing density. We tracked individual topological defects from brightfield live imaging and analyzed their trajectories and relative positions (Fig 4A). Here, the number of defects starts high at low densities and still decreasing with increasing densities, and is near constant at high densities, dropping significantly during the 30 hours gap spreading from low to high density conditions (Fig 4Bi). The distance to the nearest-neighbour of opposite charge and same charge defects increases over time after confluence, consistent with the decreasing defect density due to pair-wise annihilations (Fig 4Bii, iii).

**FIG. 4:**
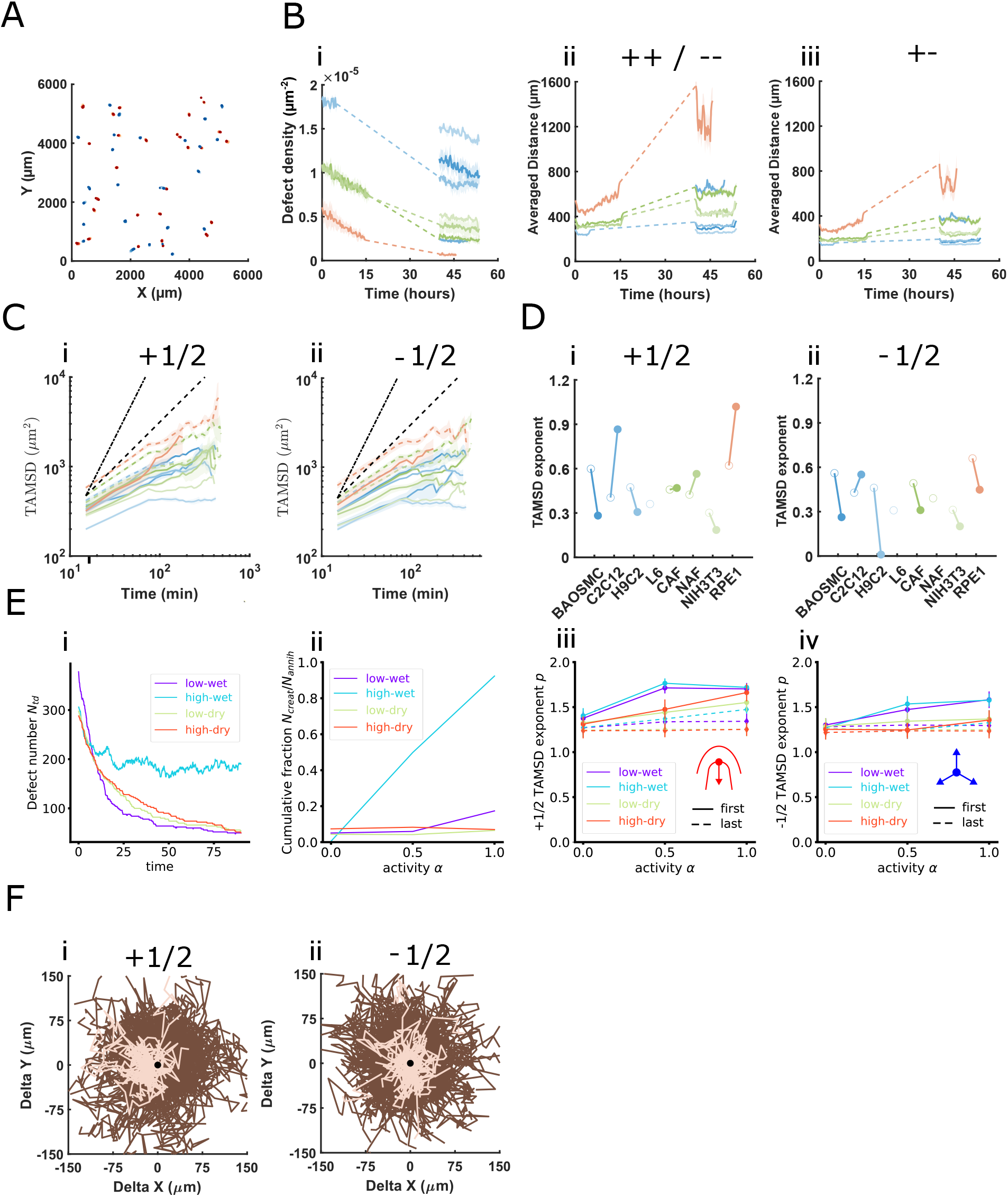
Topological defects have slow diffusive dynamics (**A**). Displacement map of +1/2 (red) and −1/2 (blue) topological defects in C2C12 monolayers at confluence, measured over a 6000 × 6000*µ*m^2^ field of view. (**B**). Analysis of topological defect dynamics: (**i**) Temporal evolution of the normalized topological defect density at confluence and 24 h after the end of acquisition, the two conditions being connected by a dotted line. (**ii**) Normalized average distance between defects of opposite topological charge. (**iii**) Average distance between defects of the same topological charge. (**C**). Time Averaged Mean squared displacement (TAMSD) analysis of (**i**) +1/2 and (**ii**) − 1/2, at confluence (dashed lines) and 2 days later (solid lines) (**D**). Slopes extracted from TAMSD distributions 48h after confluence (see methods). Open symbols correspond to the first half of the TAMSD curves, whereas filled symbols represent the second half. (**i**) +1/2 defects. (**ii**) −1/2 defects. (**E**). Theory. Evolution of the defect number as a function of time (**i**), cumulative ratio between defect creation and annihilation (**ii**), fitted exponent of the TAMSD for +1/2 defects (**iii**), fitted exponent of the TAMSD for −1/2 defects (**iv**). Exponents are computed for the first half (dashed lines) and second half (full lines) of the TAMSD curves. (**F**). Displacement maps of RPE1 (**i**) +1/2 and (**ii**) −1/2 topological defects at confluence (dark brown) and 48h after confluence (light brown). Defects displacements are aligned along the topological axis, with the tail oriented to the left.

Plotting the defects trajectories, rotated relative to the instantaneous defect orientation shows that, for most cell types and density conditions, positive and negative defect motion remains largely random and sub-diffusive, with no clear directional bias relative to the defect orientation (Fig. S5A, B). In contrast, a clear head-to-tail directed motion is observed for RPE1 +1/2 defects at confluence +48 h, consistent with the head-tail motion for a contractile active nematic expected in theory (Fig 4F).

To investigate defect motility more finely, we computed their time-averaged mean square displacement (TAMSD) (Fig. 4Ci, ii). As TAMSD are very sensitive to tracking noise induced by the discrete sampling, we compared the smallest value of the TAMSD to the square of the downsampling factor, which determines the minimum non-zero displacement (Fig. S4 A-B). For downsampling factors of 30 and 60 pixels, the TAMSD is overestimated showing that the spatial sampling is not fine enough to capture the true defect dynamics; whereas a downsampling factor of 10 pixels leads to TAMSD values well above the sampling induced threshold, excluding the effect of tracking noise on the result (Fig.S4 B). This comparison highlights the sensitivity of defect dynamics to the user-defined parameters, and the need to validate the choice of parameters based on subsequent analysis. We fitted separately the first and second half of the TAMSD, as suggested by the theoretical modeling (see after). Overall, the TAMSD are found to be sub-diffusive for nearly all cell types, at both short and long-time scales, with a typical exponent around 0.53 for positive defects and 0.38 for negative defects (Fig. 4Di, ii). One notable outlier stood out: for RPE1 at higher cell densities, the motion of +1/2 defects at long time scales becomes super-diffusive, consistent with the appearance of a directional bias in the defect trajectories (Fig. 4Fi). Overall, for the other cell types, no clear change in dynamics is observed between the short and long-time scales, and the behaviour remains predominantly sub-diffusive across all density conditions. Interestingly, reducing the cell plating density by a factor of two did not significantly affect the organization of the nematic field, nor dynamics of topological defects (Fig S3C).

For numerical results of active nematohydrodynamics, different regimes show a variety of behaviours, from passive-like annihilation of existing defects to active turbulence with equal fraction of defect pair creations and annihilations (Fig 4Eii). The TAMSD curves show a variety of dynamic regimes, with an increase in slope at a longer time scale (Fig S3D). Fitting separately the first and second half show superdiffusive TAMSD exponents, with an overall increase between short and long-time scale exponents for both positive and negative defects. For passive nematics, super-diffusion originates from a combination of defect pair annihilation forces and hydrodynamic interactions. The short-time TAMSD exponents remain independent from activity in the ‘dry’ regime, but long-time exponents are amplified by activity. This indicates that active flows enhance the mobility of defects, confirmed by the highest exponent increase in the ‘wet’ regime, Fig 4Eiii,iv. Although larger for positive defects, this active mobility effect also applies to negative defects, indicating complex collective defect dynamics. This super-diffusive behaviour is in marked contrast with experimental results, indicating an important missing ingredient to capture nematic defect dynamics.

### E. Topological defects are mechanical organisers of tissues

To explore whether the topological defects play a role in the overall tissue dynamics, we quantified the local velocity field induced by cell migration. The velocity maps were extracted from sequential brightfield images using standard Particle Imaged Velocimetry (PIV). The average velocities are on the order of a few microns per hour for all nematic cell types and climb up to 10-15 *µ*m/h for the isotropic epithelial monolayers (HBEC and MDCK) (Fig 5A).

**FIG. 5:**
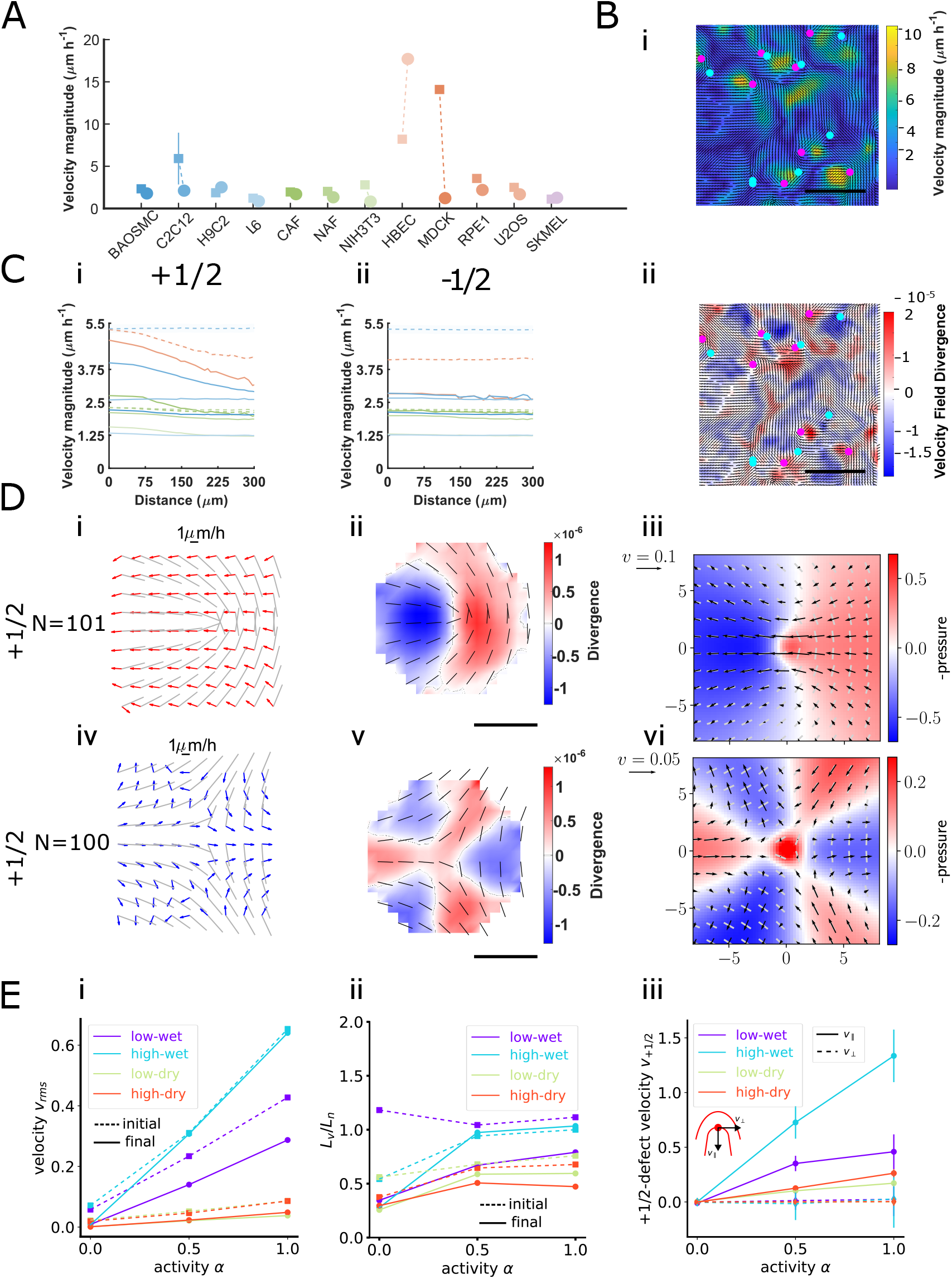
Topological defects are mechanical organisers of tissues (**A**). Average cell velocity magnitude measured at confluence (squares) and 2 days after confluence (circles) using particle image velocimetry (PIV) for all cell types analyzed. (**B**). Velocity and divergence fields in RPE1 monolayers imaged 2 days after confluence. (**i**) Magnitude of the velocity field displayed as a color map. Positive (+1/2) topological defects are shown in magenta and negative (−1/2) defects in cyan. Scale bar, 500*µ*m. (**ii**) Divergence of the velocity field displayed as a color map, computed from the same velocity data. (**C**). Radial distribution of the average cell velocity magnitude around (**i**) +1/2 and (**ii**) −1/2 topological defects, with distance measured from the defect center, at confluence (dashed lines) and 2 days post confluence (solid lines). (**D**). Analysis of velocity fluxes around topological defects from NIH3T3. (**i**) Averaged +1/2 defect configuration showing the nematic orientation field (grey) and the averaged velocity field (red). Spatial averaging is performed using nodes of size 30 pixels (19.5*µ*m). (**ii**) Corresponding averaged divergence map of the velocity field around +1/2 defects. (**iii**) Theoretical divergence map around +1/2 defects, superposed with director field (white) and velocity field (black) in the ‘low-dry’ regime at *α* = 0.5. (**iv-vi**) Same analyses as in (**i-iii**), respectively, for −1/2 defects. (**E**). Theory. (**i**) Average tissue velocity as a function of activity.(**ii**) Ratio between nematic and velocity correlation length as a function of activity. (**iii**) Average tissue velocity at +1/2 defect positions as a function of activity.

The velocities are heterogeneous, with large spatial fluctuations, where higher magnitudes seem to correlate with topological defects position (Fig 5B). Similarly, on average, velocities are not correlated with the local orientation as velocities with small amplitude have random orientation. Instead, restricting the analysis to the top 5% of velocities with the largest amplitudes shows a clear positive correlation (Fig. S5B). Plotting the velocity magnitude as a function of distance to defect core reveals that only positive half defects are associated with local maxima of the velocity (Fig 5C, S5A). In addition to peaking, the velocities are spatially correlated at positive topological defects: the cellular flows are oriented from head to tail for all cell types (Fig 5Di-ii, Fig S5 C-D). For negative defects, the flows follow the same pattern: they are oriented outward the three bends and inwards along the three branches, leading to small average flows at the defect core (Fig 5C ii, Fig S5A i ii). Both match the expected active flows for a contractile nematic around defects, Fig 5E.

The flow divergence also shows significant fluctuations at the tissue scale (Fig 5 Bii) and a stereotypic pattern around topological defects: convergent at the tail of the +1/2 and divergent at the head. This pattern is consistent with the theoretical predictions around a +1/2 defects in a contractile system with pressure-dependent flow divergence due to cell proliferation [37]. Similarly the pattern around −1/2 defects matches the predictions [37]. In incompressible active nematics, for a sufficiently slow proliferation rate, velocity divergence is controlled by the pressure map around defects generated by active stress gradients (Sec. IV.R)). Hence even without proliferation, the same patterns are observed in numerical results with incompressibility (Fig 5D iii, vi). Measuring the cell density from nuclei stains around topological defects does not show any accumulation or depletion (Fig. S5E). This suggests that these convergent/divergent flows do not translate into long lasting particle density variations, possibly due to the highly motile nature of the cells which could relax density gradients.

In numerical results, the average velocity also tends to decrease over time, with large dependence on the different regimes (Fig 5Ei). Since the theory assumes material flows to be generated by gradients in orientation, the velocity depends on the defect density in a precise way, with scaling exponents expected for different regimes (Fig S5 G). It also implies the velocity correlation length *L*_*v*_ to be influenced by the nematic correlation length *L*_*n*_. Numerical results show that *L*_*v*_ is essentially lower than *L*_*n*_ (Fig 5Eii, S5F). This is in agreement with experimental results on Fig. S5F, indicating a dry regime with strong substrate friction where flows are controlled by the hydrodynamic length *l*_*h*_. The material velocity at +1/2 defects is clearly directed towards the tail and correlates with activity amplitude (Fig 5Eiii), in agreement with contractile active stress and experimental findings (Fig 5Di).

## III. DISCUSSION

By systematically exploring the impact of the various user-defined parameters in the nematic field analysis, we reveal the sensitivity of the measured static and dynamic properties, underscoring the importance of reporting these values in future publications. We propose guidelines to select these parameters in a data-informed manner and further improve comparability of reported data.

Deploying this extensive analysis on twelve different cell types of various origins uncovers a clear distinction between isotropic and nematic cellular tissues. Among nematic tissues, most nematic properties (correlation length, defect density and distribution, splay to bend ratio, defect velocities and MSD exponents) are of similar values, underlining the universality of the liquid crystal nature across living tissues. Nevertheless, all these properties are highly time-dependent, possibly due to an interplay with cell density, further emphasizing the need for robustness in analysis and reporting to enable cross-project comparisons. We further combine the nematic analysis with PIV analysis to measure local cellular flows. Topological defects are hotspots of the velocity field within mostly frozen tissues, with cells streaming at the defect core. Interestingly, all the cell types analyzed here exhibit directional cellular flows at the defect core, oriented head-to-tail, compatible with contractile active nematics. Yet when confined in channels [5, 8, 9], cell types like C2C12 or RPE1 behave more as extensile nematics. Although investigated in [28, 29] the question of the activity sign of cell assemblies remains an open question.

In the literature, defect motion is usually inferred from the cellular flow patterns around the defect core, consistent with the expected properties of active nematics. This leads to the assumption of ballistic motion for +1/2 defects. To analyse this, we performed a direct single defect tracking analysis, reporting defect trajectories and TAMSD analysis, which revealed surprising defect dynamics. While the flow patterns at the defect core are clearly directional, oriented head-to-tail as found in an active contractile nematic fluid, the defect motion is diffusive and not directional. Our findings show that a double characterization of individual defect dynamics and cell material flows is essential to make a faithful comparison to theory. We show that the hydrodynamic theory of active nematics captures static nematic properties and material flows around defects. However, for all regimes considered, it fails to explain the subdiffusive dynamics of individual defect trajectories.

In fact, the self-propulsion of +1/2 defects (ballistic and directional) is predicted by theory for isolated conditions, a single defect in an infinite nematic environment [37, 38]. In our numerical results, the long-range defect-defect interactions mediated by nematic elasticity induce rotational diffusion of +1/2 defects and prevent perfect ballistic dynamics. In parallel, defect pair annihilation induces super-diffusive motion even in the passive case, underscoring the importance of collective effects from defect interactions to interpret data from defect trajectories. Yet even collective effects fail to predict the subdiffusivity found in experiments.

As proposed recently [21, 30], the extracellular matrix deposited by cells could act as a freezing medium for the nematic texture and prevent persistent defect motion. An alternative hypothesis is the increasing density due to continuous proliferation [10, 23]. Although our active nematic theory assumed two-dimensional incompressibility, hence a vanishing flow divergence, experimental cell flows are not divergence-free and exhibit coherent patterns around topological defects. The theory could be extended with a cell density field to account for compressibility. Then, density-dependent parameters could introduce aging dynamics and contribute to the sub-diffusive dynamics of defect trajectories.

Overall, our work establishes a standardized framework for the characterization of nematic order in cellular tissues, enabling reproducible analyses and facilitating broader adoption by the biological community. By systematically comparing multiple cell types, we identify robust ranges for key nematic observables, providing experimentally grounded benchmarks for theoretical modeling. At the same time, the observed subdiffusive defect dynamics highlight a fundamental limitation of current active nematic theories in describing living systems. These results call for extended theoretical frameworks incorporating additional ingredients such as density variations, extracellular matrix interactions, or aging effects. More broadly, our study bridges experimental practice and theory, and sets the stage for a quantitative understanding of active nematics in living matter.

## IV. MATERIAL AND METHODS

### A. Cell culture

Except if indicated otherwise, all cells were cultured in T75 tissue culture treated flasks (Corning REF 430641U) using DMEM (Lonza REF (BE)12-733) supplemented with 10% FBS and 1% Penicillin/Streptomycin (Gibco 15140122) prior to plating after one or two passages at 1 ×10^4^ to 1 ×10^5^ cells/mL on tissue culture treated plastic 6 well plates (Corning) at two different seeding densities and with 3 wells per conditions. RPE1 cells were cultivated using DMEM/F-12 (Gibco). hBEC cells were cultivated using Bronchial Epithelial Cell Growth Medium (BEGM-Lonza CC-3170). U20S were cultivated using DMEM GlutaMAX medium (Gibco, REF 61965-026). SKMEL were cultivated using EMEM (Gibco). All cell lines were tested for mycoplasma contamination using Mycoplasmacheck PCR Detection.

### B. Live Nematic field imaging

To follow the evolution of the nematic fields along experiments a daily acquisition of a 7 ×7 grid of images was performed (14000 ×14000 *µ*m^2^, with 200 *µ*m overlap between each image on Molecular Devices ImageXpress Micro C microscope, Nikon 10× phase contrast objective, Ph1 ring, with a pixel size of 0.635 *µ*m), at the center of each well over the course of a week. To follow the dynamics of the tissues, an overnight acquisition of a 5 ×5 grid of images was performed around confluence and two days after confluence (10000 ×10000 *µ*m^2^, with 200 *µ*m overlap between each image), at the center of each well with one image every fifteen minutes.

### C. Immunostaining and fluorescence imaging

Prior to imaging cells were fixed by adding 200 *µ*L of 4% PFA in PBS for thirty minutes in each well. Cells were then permeabilized using 500 *µ*L of 0.1% Triton X in PBS and stained for nuclei (Hoechst 33342 1:6000 in PBS) and actin filaments (Phalloidin 488 nm 1:400 in PBS). Acquisition of stained samples was performed on a Molecular Devices ImageXpress Micro C microscope using a 10× plan Apochromat objective. A 7 × 7 grid of images (14000 ×14000 *µ*m^2^, with 200 *µ*m overlap between each image) was also captured at the centre of each well.

### D. Image processing

All analyses were performed using custom MATLAB codes developed or adapted in the laboratory. In total, 12 cell types were investigated in this study: BAOSMC, C2C12, H9C2, and L6 (myoblasts); CAF, NAF, and NIH3T3 (fibroblasts); HBEC, MDCK, RPE1, and U2OS (epithelial cells); and SKMEL. Among them, eight displayed nematic cell tissue organization, namely BAOSMC, C2C12, CAF, H9C2, L6, NIH3T3, NAF, and RPE1. Quantitative analyses comparing confluence and two days post confluence were restricted to four cell types (CAF, NAF, H9C2, and RPE1) for which three independent fully confluent biological replicates were available.

### E. Orientation field extraction

Cellular and actin orientation were obtained using OrientationJ, an ImageJ plugin developed by the EPFL Bio-Imaging and Optics Platform. Orientation is obtained by computing the gradient of intensity *I*(*x, y*) for each pixel localized at position (*x, y*) over a defined distance (feature size), yielding the spatial derivatives *I*_*x*_ = ∂*I/*∂*x* and *I*_*y*_ = ∂*I/*∂*y*. From these quantities, a structure tensor **J** is computed for each pixel as

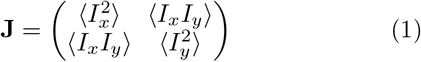

which can be decomposed into isotropic and nematic components

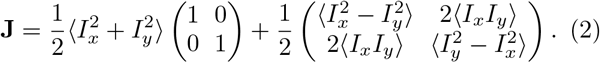

Here ⟨ · ⟩ denotes local averaging using a Gaussian window whose size is defined by the feature size. The principal angle of orientation *θ* is extracted from the nematic part of the structure tensor, whereas the local correlation of orientation between neighboring pixels (coherency) is extracted from the eigenvalues *λ*_1_ ≥ *λ*_2_ of **J**. The orientation angle is given by

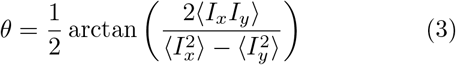

and the coherency is defined as

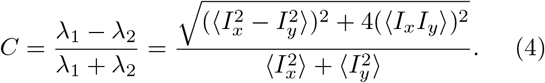

This quantity varies from 0 (isotropic structure) to 1 (perfectly aligned structure).

A discretization of orientation and coherency is performed by averaging pixels inside grids of user-defined size (downsampling size) (see Fig 1D). The averaged orientation for each position in the grid is saved in a result table (CSV file).

### F. Nematic correlation length measurement

The nematic correlation function *C*_*nn*_ was used to measure the range over which orientation propagates in the nematic field.

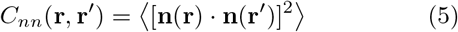

for a director field **n**(**r**) = cos *θ*(**r**) **e**_*x*_ + sin *θ*(**r**) **e**_*y*_ and a position vector **r** = *x* **e**_*x*_ + *y* **e**_*y*_. The two-point correlation function was then transformed into a radial function with 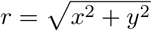, showing an exponential decay distribution.

The value of *r* in microns at which the exponential reached *e*^−1^ was defined as the correlation length. The correlation length was systematically averaged for each cell type on three 14000 × 14000 *µ*m^2^ images and at each time point captured during live imaging on 5590 × 5590 *µ*m^2^ images.

### G. Nematic order parameter computation

The nematic order parameter was computed in each position of the director field using

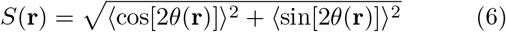

where ⟨·⟩ indicates the average of director angles *θ* over a circular region in the position matrix. In this study, except if stated otherwise, order was computer on a radius of 1 down-sampling factor size (10 pixels).

### H. Topological defect detection

For each candidate point located in regions where the nematic order parameter is below a defined threshold (referred as S Threshold, value 0.5), orientation differences are computed sequentially between neighbouring points and summed. The total rotation divided by 2*π* identifies the topological charge. A resulting value of +0.5 identifies a +1/2 defect whereas −0.5 identifies a −1/2 defect.

Due to the finite spatial resolution of the grid and local noise in the orientational field, a single physical defect can be detected at several neighboring nodes, forming a small cluster of nodes carrying identical charges. To avoid multiple detections of the same defect, nodes of identical sign located within a distance smaller than four grid spacings were merged. The position of the resulting defect was defined as the center of mass of the clustered nodes, and the defect charge was assigned as ±1/2 according to the sign of the average charge within the cluster.

### I. Defect tracking

Topological defect tracking was performed using a global optimal assignment algorithm applied to consecutive matrices of defect positions. The analysis was carried out on time-lapse movies acquired at confluence and from 24 h to 48 h after confluence. Movies consisted of *N* = 3 fields of view with an image size of 8600 × 8600 pixels, acquired every 15 min over a total duration ranging from 6 h to 15 h. At each time point, +1/2 and −1/2 defects were identified independently from the topological charge matrix.

Defect candidates were then linked across frames using a distance-based cost matrix and a global Hungarian assignment. A maximal linking radius of 10 grid nodes (100 pixels) was imposed to prevent unphysical associations. Defects were allowed to persist without observation for up to 5 consecutive frames to account for intermittent detection failures. Defects with lifetimes shorter than three time steps were excluded from further analysis.

### J. Time-averaged mean squared displacement (TAMSD)

For each defect trajectory, the time-averaged mean squared displacement (TAMSD) was computed. For a trajectory **r**(*t*) where *t* is a frame number, the squared displacement is Δr^2^(t, Δt) = |**r**(*t* + Δt) −**r**(*t*) |^2^. The TAMSD is computed as

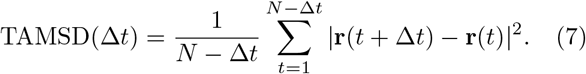

Time lags were restricted to Δt ≤N/2. Squared displacements *s* were converted to physical units using *s* = Pixel size × Downsampling factor.

#### K. Extraction of TAMSD scaling exponents

The TAMSD was first computed individually for each defect trajectory. For each experimental condition (cell type and density), the TAMSD curves obtained from all tracked defects were then averaged to obtain a mean TAMSD curve ⟨TAMSD(Δt) ⟩. The MSD scaling exponent *α* was extracted from this mean TAMSD curve. Defect dynamics were characterized by fitting the TAMSD to a power-law relationship TAMSD(Δt) ~ (Δt)^α^. Linear regression was performed on the logarithmically transformed data log(TAMSD) = *α* log(Δt) + *b*. Fits were performed only when at least two valid MSD points were available within the fitting window.

### L. Instantaneous defect velocity

Instantaneous velocities were computed from frame-to-frame displacements along each defect trajectory. For consecutive frames, the displacement components were defined as *d*_*x*_(*t*) = *x*(*t* + 1) −*x*(*t*) and *d*_*y*_(*t*) = *y*(*t* + 1) −*y*(*t*). After conversion to micrometers *d*_*x*_[*µ*m] = *d*_*x*_ *s, d*_*y*_[*µ*m] = *d*_*y*_ *s*, the instantaneous displacement magnitude was calculated as

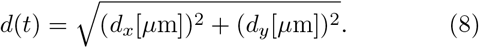

Given the imaging interval of 15 minutes (0.25 h), instantaneous velocity was computed as *v*(*t*) = *d*(*t*)/0.25, resulting in velocities expressed in ·*µ*m h^−1^.

The mean velocity of each defect was obtained by averaging instantaneous velocities over all valid time steps of the trajectory. Defects velocities extracted from the three replicates were then averaged together for each condition of cell density.

### M. Directional displacement relative to defect orientation

To analyse defect motion relative to their intrinsic polarity, defect trajectories were realigned according to their polarity. For +1/2 defects, the reference direction was defined by the orientation of the comet tail. For −1/2 defects, trajectories were aligned relative to the orientation of the upper arm of the trefoil structure. Trajectories were rotated so that this reference direction defines a common axis, enabling the analysis of defect motion relative to their intrinsic orientation.

### N. Frank constant ratio computation

Frank constant ratios were computed based on the method developed by Zhang et al. [35]. Square windows of 15 ×15 vectors centered on +1/2 topological defects were rotated to align the tail of the defect (determined during defect detection) to the left. The horizontal axis from the center of the defect to the right was set as the initial rotation line of the polar angle *ϕ. θ* was set as the local angle of the director in the polar coordinates basis, corresponding to a rotation of *ϕ* to the position of a director **n** in the matrix of positions. For each cell type, a combination of at least 10 images (5000 ×5000 *µ*m^2^) was analyzed and averaged. Theoretical defects of known Frank constant ratios were generated as calibration curves to assess the experimental distributions.

### O. Cell flow measurement and analysis around topological defects

Cellular flow fields were quantified from time-lapse image series acquired every 15 minutes using the PIVlab Matlab toolbox [39]. Particle Image Velocimetry (PIV) analysis was performed using four iterative passes with progressively decreasing interrogation window sizes (from 450 ×450 *µ*m^2^ to 150 ×150 *µ*m^2^) and a 50% overlap between adjacent windows. All analyses of cellular dynamics were performed over the entire duration of the acquisitions (minimum 6 hours), corresponding to 5× 5 tiled images (8600 ×8600 px^2^), at confluence and 48 hours post-confluence. Analyses were conducted on all detected topological defects across all time points and replicates. To characterize cell flows around topological defects, divergence fields were computed from the velocity matrices in the vicinity of each defect. Defects were rotated such that the defect tail was systematically oriented toward the left, and the associated divergence matrices were rotated accordingly. Divergence values were then averaged over all defects and all replicates for each cell type. An equivalent procedure was applied to velocity fields, including both velocity magnitude and orientation. Radial profiles of velocity magnitude were obtained by extracting the velocity magnitude as a function of radial distance from the defect core, followed by averaging over all defects and replicates.

### P. Nuclei intensity around topological defects

For the analysis of nuclear intensity and cytoskeletal organization, cells were stained with DAPI to label nuclei and with Phalloidin-488 to visualize actin filaments. Images were acquired using a Nikon 10× objective as 7 ×7 tiled fields of view (11600 ×11600 ×px^2^), with three replicates per cell type. Nuclear intensity was extracted from the DAPI channel, while actin orientation fields were computed from the phalloidin signal. Detected topological defects from actin orientation fields were rotated to align their tails toward the left, and the corresponding intensity maps were rotated accordingly. Averaged distributions of nuclear intensity and actin orientation were then computed by pooling all defects across all replicates for each cell type.

### Q. Velocity correlation length measurement

The spatial extent of collective migration was quantified using the normalized velocity–velocity correlation function

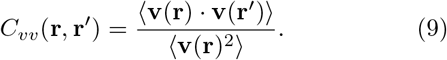

where **v**(**r**) = *v*_*x*_(**r**)**e**_*x*_+*v*_*y*_(**r**)**e**_*y*_. Prior to correlation analysis, the mean drift was subtracted from both velocity components to remove global translation of the monolayer. The two-dimensional correlation map was computed using a Fourier-based approach and subsequently averaged over the radial coordinate 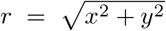 to obtain a one-dimensional profile exhibiting an approximately exponential decay. The correlation length *L*_*v*_ was defined as the distance (in *µ*m) at which the normalized correlation function decayed to *e*^−1^ of its value at the origin.

### R. Theoretical description

Nematohydrodynamic equations for the nematic **Q**(**r**, *t*) and velocity **v**(**r**, *t*) fields are solved under periodic boundary conditions in a two-dimensional rectangular box. In two dimensions, the nematic tensor **Q** can be decomposed into an order parameter *S* and a director **n**, such that **Q** = *S*(2**n**⊗ **n** −**I**), where **I** is the identity tensor. We assume two-dimensional incompressibility for the velocity field, corresponding to the equation ∇ · **v** = 0. The nematic free energy is written as

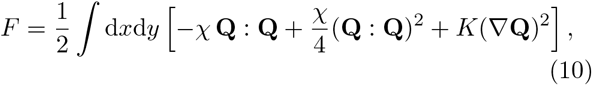

where *χ* is the ordering parameter and *K* the orientational elastic parameter. From this free energy, one can derive the molecular field **H** = *χ*(1 −*S*^2^)**Q** + *K*Δ**Q** that represents equilibrium torques. The dynamics of the nematic tensor then follows as

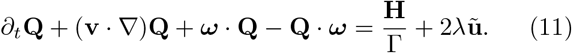

The left-hand side represents the co-rotational derivative with a rotational viscosity Γ and vorticity 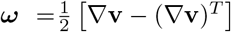. On the right-hand side, *λ* is the flow-alignment parameter that couples nematic dynamics to the strain rate 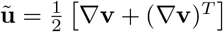.

Force balance for a two-dimensional fluid in contact with a substrate reads

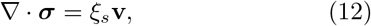

where ***σ*** is the total tissue stress and *ξ*_*s*_ is the substrate friction parameter. The stress tensor is decomposed into pressure, viscous stress with viscosity *η*, Ericksen stress ***σ***^*E*^ = *K*∇ **Q** ⊗ ∇**Q**, reactive stress and active stress with coefficient *α*:

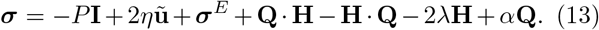

To make equations dimensionless, one uses the nematic core length 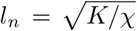 as a length scale, the nematic relaxation time *t*_*n*_ = Γ*/χ* as a time scale, and the ordering parameter *χ* as a stress scale. Four dimensionless parameters remain, *λ, η, α, ξ*_*s*_, and the same notation is kept. Viscosity is fixed at a value *η* = 1.5, and the four regimes are defined with the following doublets (*λ, ξ*_*s*_): (0, 0) for ‘high-wet’, (1.1, 0) for ‘low-wet’, (0, 1.5) for ‘high-dry’ and (1.1, 1.5) for ‘low-dry’.

The partial differential equations are solved using a Fourier spectral method on a regular grid with node spacing Δx = Δy = 0.25 and time step Δt = 10^−4^. A system size of *L* = 256 is used, with *N* = 1024 nodes per dimension, and a total simulation time *t*_sim_ = 100, or *N*_*t*_ = 10^6^ number of time steps. Nematic, velocity and pressure fields are saved every 1000 time steps.

Linear stability analysis is done around a uniform state with horizontal nematic order, by linearizing the equations and performing a spatial Fourier transform. For perturbation along (bend) or perpendicular (splay) to the nematic orientation, an instability emerges for a critical activity

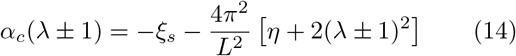

where the sign + (−) correspond to bend (splay). The wavenumber associated to the most unstable growth-rate is found numerically in the parametric plane (*α, λ*) without and with substrate friction, Fig. 1Gi.

To quantify spatial quantities around single defects, squares of size 16 ×16 are rotated to align defect orientations and averaged over the last 50 consecutive frames, from *t* = 95 to *t* = 100. For the shape anisotropy of +1/2 defects, the difference of the orientation profile *θ*(*ϕ*) to the case *K*_1_ = *K*_3_ (*θ*(*ϕ*) = *ϕ/*2) is computed numerically for each defect and integrated from *ϕ* = 0 to *ϕ* = *π*. The result for perfect-splay or prefect-bend is equal to ±*π*^2^/8, and the factor *ϵ* is obtained by normalization with this quantity (Fig. 3Civ).

To generate flow divergence maps around defects, we start from the equation of tissue proliferation under 2D incompressibility ∇ ·**v** =− *κ*(*P* −*P*_*h*_), where *P* is the pressure, *κ* a coupling parameter measuring the pressure-dependent proliferation, and *P*_*h*_ the homeostatic pressure parameter. As mentioned in the main text, a small coupling constant generates non-zero flow divergence with negligible change of the flow dynamics generated by active stress gradients. In those conditions, the flow divergence map around defects should be equivalent to the opposite of the pressure map, −*P*, which is computed in Fig. 5D.

## ACKNOWLEDGMENTS

We thank A. Babataheri, P. Guillamat, and D. Vignjevic for providing, respectively, the hBEC and BAOSMC, NIH3T3, and CAF and NAF cell lines used in this study. We acknowledge the ACCESS Platform at UNIGE for discussions and support with microscopy acquisitions. We also thank P. Guillamat for insightful discussions and assistance with the analysis code.

AR acknowledges funding from the Swiss National Fund for Research grant numbers #CRSII5 189996 and #310030 200793 and the European Research Council Synergy grant number #951324-R2-TENSION.

## AUTHOR CONTRIBUTIONS

N.R. contributed to methodology, investigation, formal analysis, visualization and writing—review and editing. M.D. contributed to conceptualization, formal analysis, investigation, and writing of the original draft. A.R. contributed to conceptualization, funding acquisition, project administration, and writing—review and editing. C.D. contributed to conceptualization, supervision, and writing of the original draft.

## V. SUPPLEMENTARY FIGURES

**FIG. S0:**
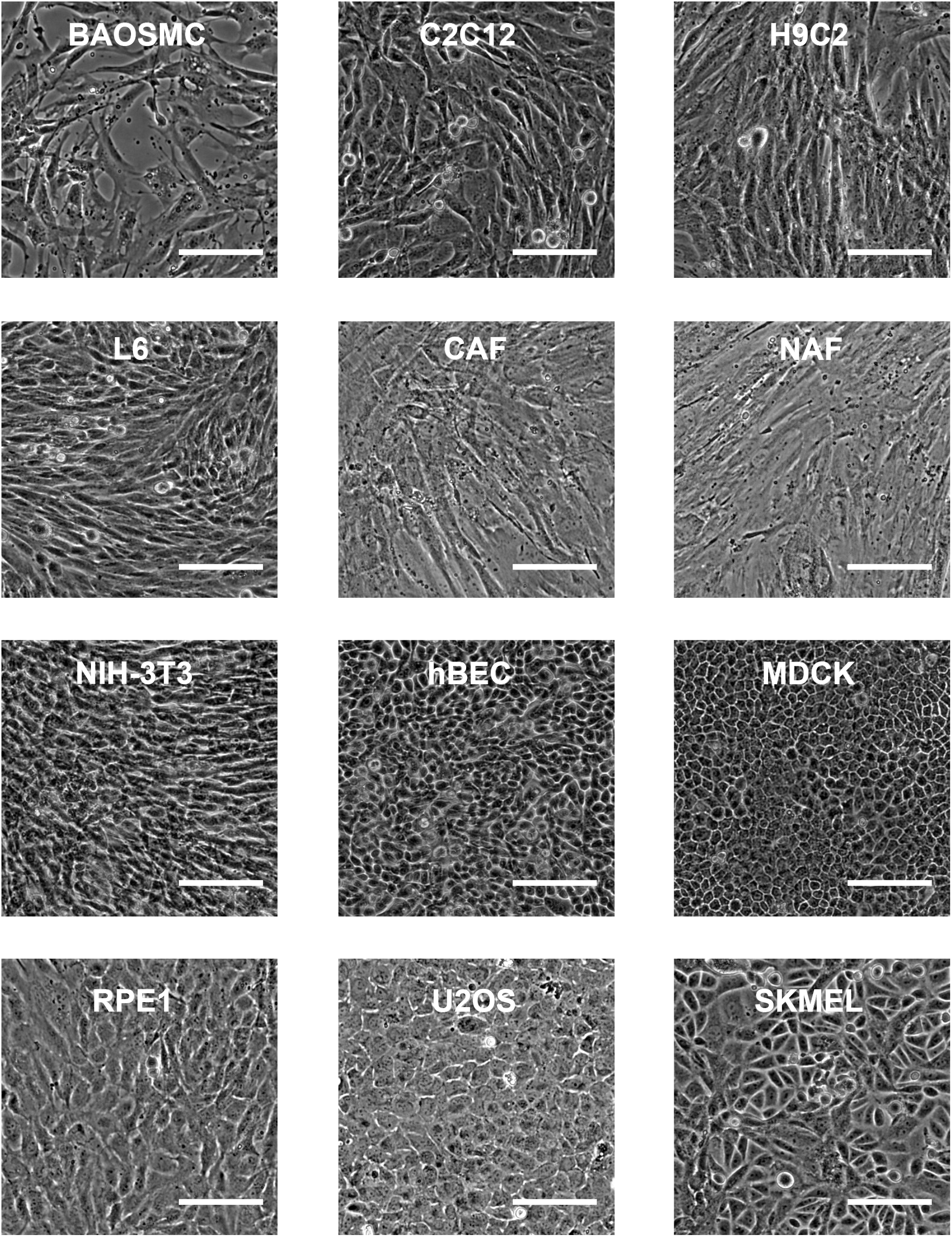
10X Phase contrast Brightfield images of the twelve studied cell types (NAF and CAF were captured with 10X DIC). Scale bar represents 100 *µm*

**FIG. S1:**
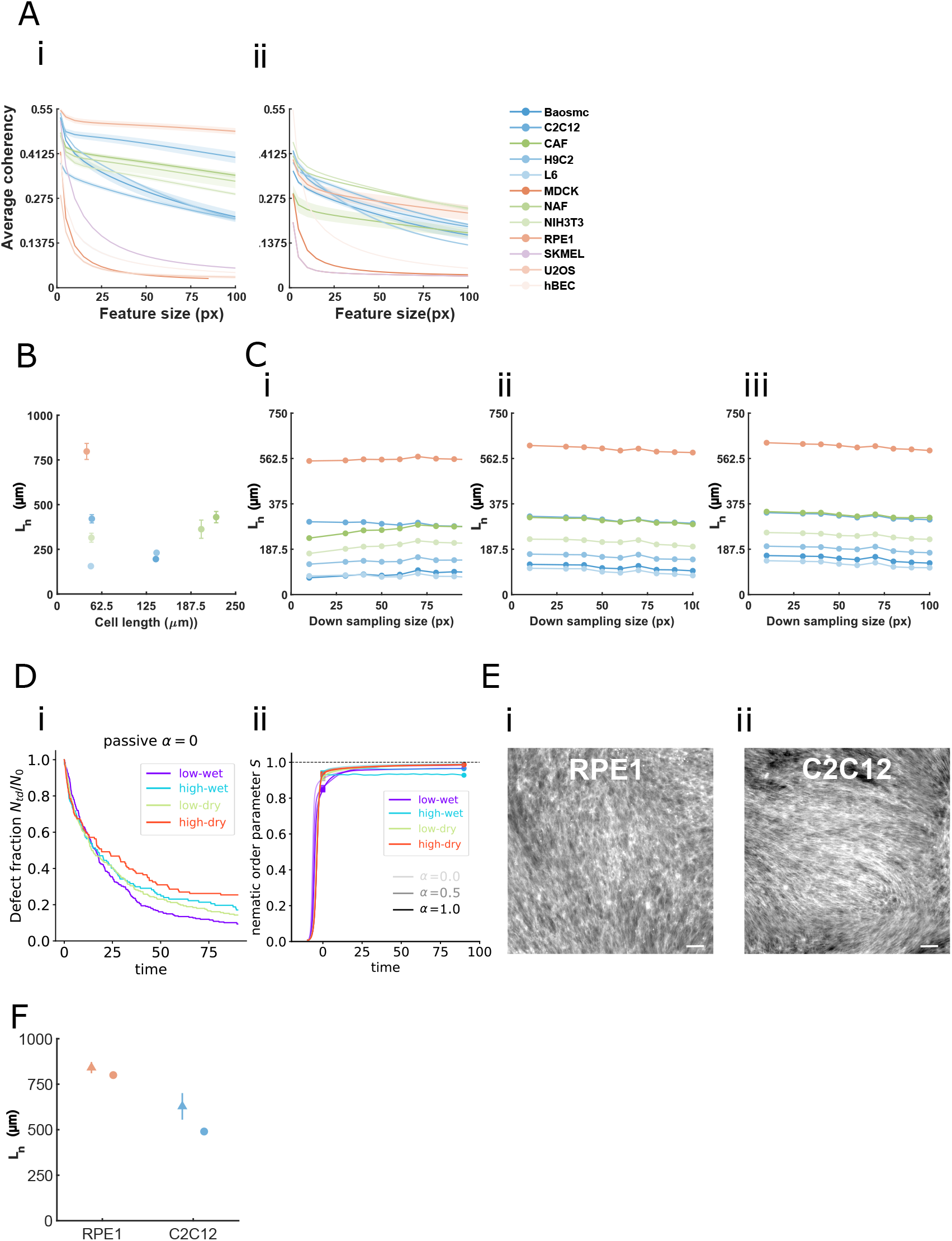
Measurements were performed on *N* = 3 images per condition. (**A**). Distribution of the average orientational coherency as a function of the feature size for all cell types analyzed (**i**) at confluence, (**i**) two days after confluence. (**B**). Distribution of the average correlation length (*µ*m) as a function of cell size for all cell types analyzed, 48h after confluence. (**C**). Distribution of the average correlation length as a function of the OrientationJ analysis down sampling factor for all cell types analyzed at confluence. Feature size (**i**) 5 pixels, (**ii**) 25 pixels (**iii**) 50 pixels. (**D**) Evolution of the defect fraction for the different regimes of a passive system (**i**), and evolution of the average nematic order parameter *S* for different regimes and activities (**ii**). (**E**) Confocal image of Phalloidin-GFP stained actin,(**i**)RPE1 (**ii**) C2C12, scale bar represent 100(*µ*m). (**F**) Nematic correlation length of actin orientation field (triangle) and measured on brightfield images 48h after confluence (circle).

**FIG. S2:**
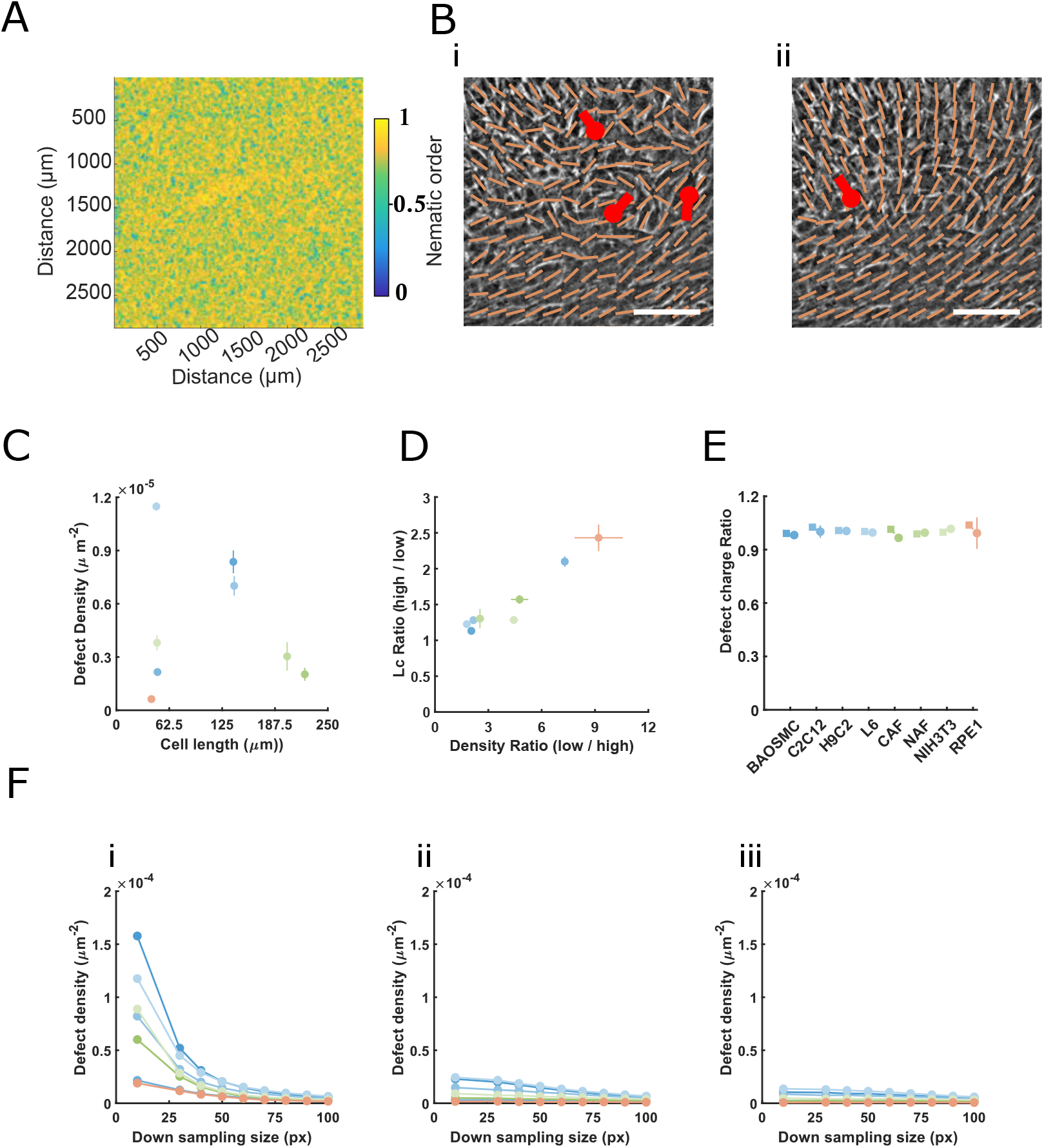
Nematic order maps and topological defect statistics across cell types and analysis parameters. (**A**), Nematic order color map of MDCK monolayers computed using Equation 2 with a neighbouring distance of 60 pixels. (**B**), Topological defect detection in RPE1 monolayers, scale bar is 100*µ*m. (**i**) Detected +1/2 topological defects overlaid on orientation fields computed from the images shown in Figure 1, using an orientation-analysis window size of 2 pixels. (**ii**) Same images analyzed using an orientation-analysis window size of 50 pixels. (**C**), Average topological defect density measured 2 days after confluence plotted as a function of cell length. (**D**), Relationship between changes in topological defect density and changes in correlation length between confluence and 2 days after confluence. The ratio of defect density at confluence to defect density 2 days after confluence is plotted as a function of the ratio of correlation length at confluence to correlation length 2 days after confluence.(**E**), Ratio of positive/negative topological defects at confluence (square) and two days after confluence (circle). (**F**), Distribution of the average defect density as a function of the OrientationJ analysis down sampling factor for all cell types analyzed at confluence. Measurements were performed on *N* = 3 images per condition with one image size of 11, 600 × 11, 600 pixels (1 pixel= 0.65*µ*m). Feature size (**i**) 5 pixels, (**ii**) 25 pixels (**iii**) 50 pixels.

**FIG. S3:**
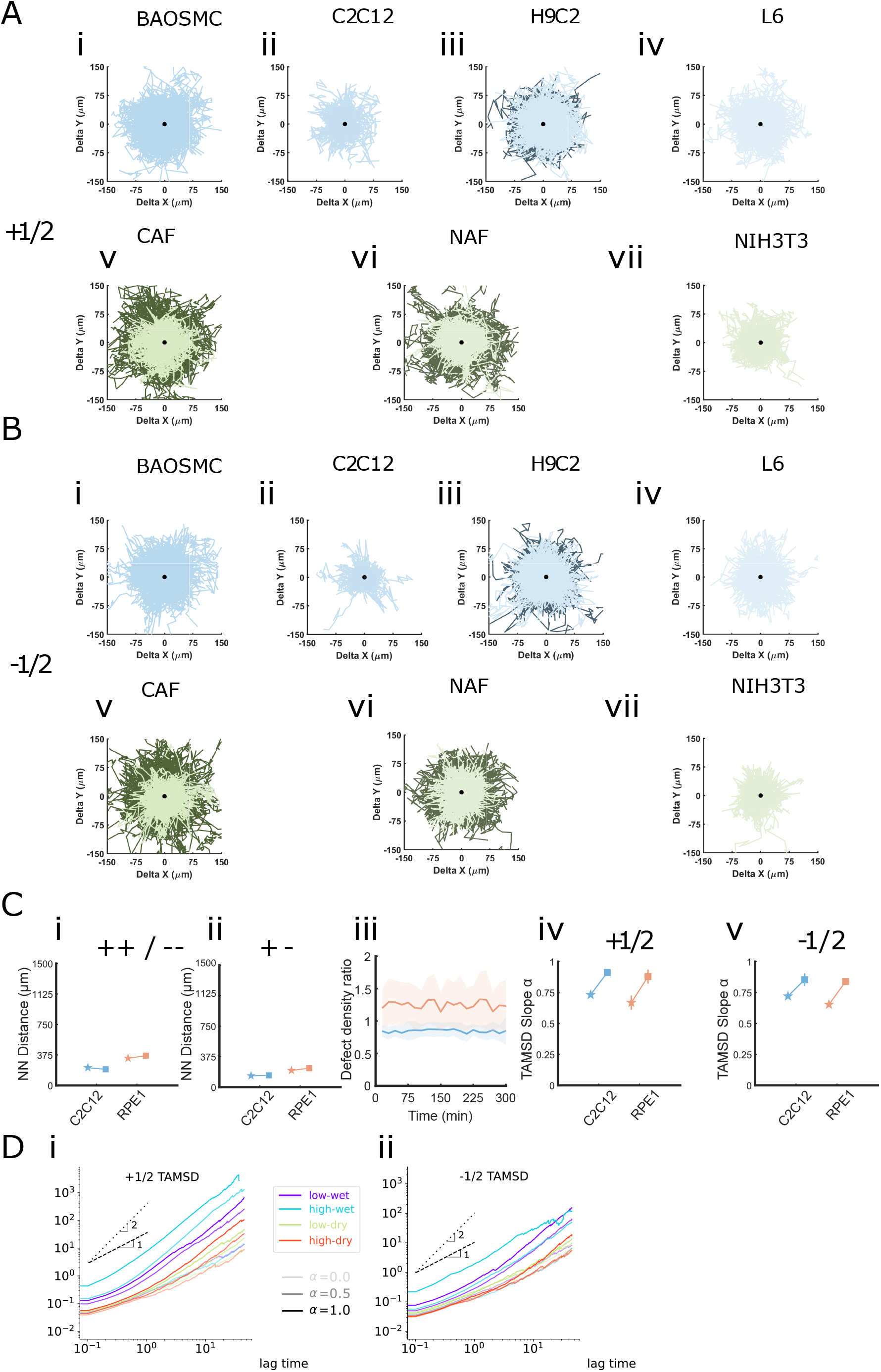
Displacement of topological defects in monolayers. (**A-B**). Displacement maps of +1/2 (**A**) and −1/2 (**B**) defects aligned on topological axis (tail to the left, or trefoil up) at confluence (dark colour) and 48h after confluence (light colour) (**i**)BAOSMC, (**ii**) C2C12,(**iii**) H9C2, (**iv**)L6, (**v**) CAF, (**vi**) NAF, (**vii**) NIH3T3. (**C**). Analysis of C2C12 and RPE1 topological defect dynamics when cells were seeded at lower density(star) and high seeding density (square). Average nearest-neighbour (NN) distance (**i**) between defects of the same topological charge, (**ii**) between defects of opposite topological charge (**iii**) Temporal evolution of topological defect density ratios (high/low) at confluence between low and high seeding condition. Average Slopes extracted from MSD distributions. (**iv**) +1/2 defects. (**v**) −1/2 defects. (**D**). Theoretical TAMSD curves for the different parametric regimes, for +1/2 (i) and −1/2 (**ii**) defects. (**E**). Ratio of Nearest Neighbour Distance to Correlation length 48h after confluence.

**FIG. S4:**
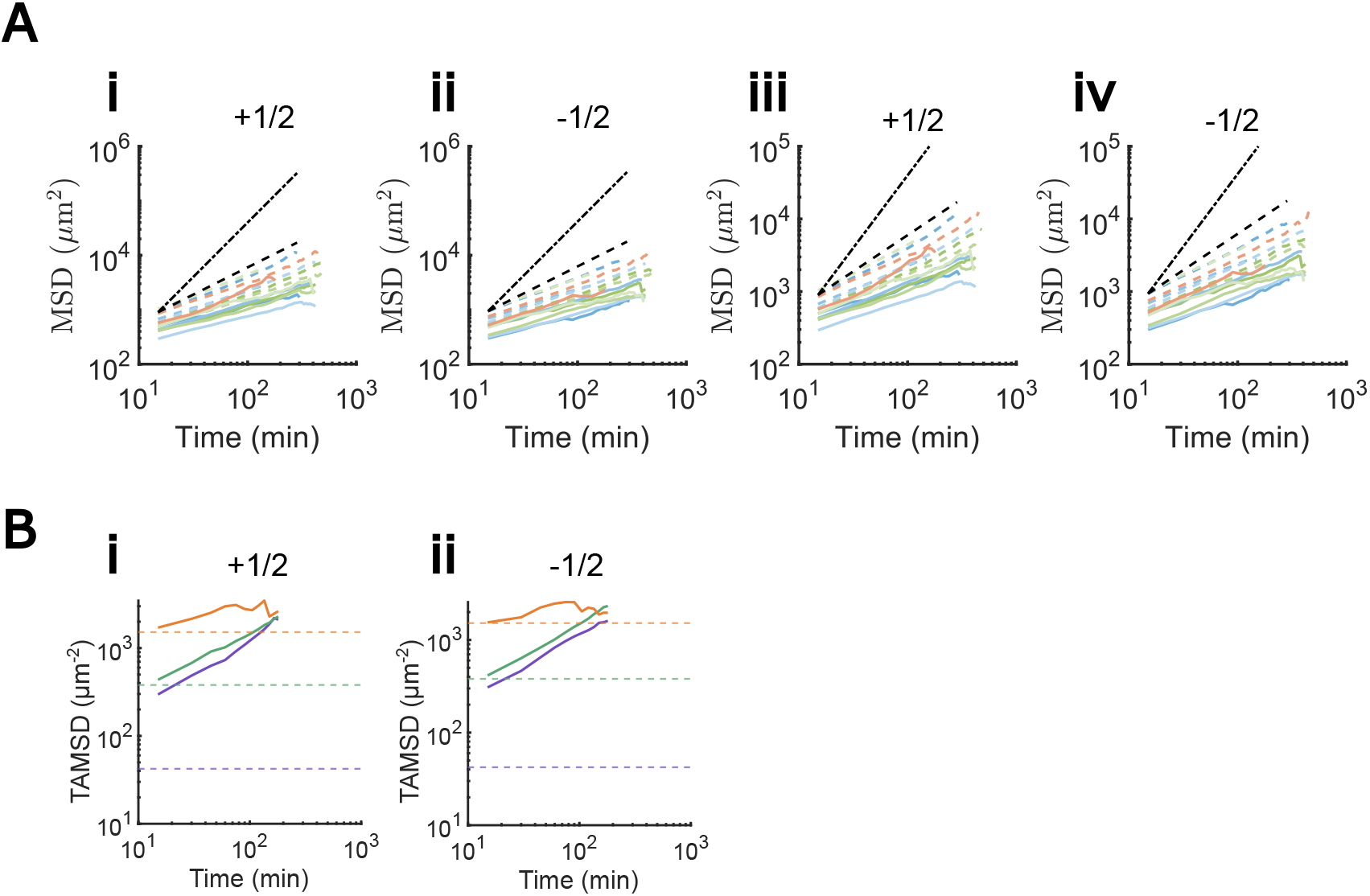
(**A**). Time Averaged Mean squared displacement (TAMSD) analysis with a down sampling size of 30 pixels (**i**) +1/2 and (**ii**) −1/2, and 60 pixels **iii**) +1/2 and (**iv**) at confluence (dashed lines) and 2 days later (solid lines). (**B**) RPE1 Time Averaged Mean squared displacement (TAMSD) analysis two days after confluence with a down sampling size of 10 pixels (violet), 30 pixels (green), 60 pixels (orange). Dashed lines represent the square of the down sampling size, for (**i**) +1/2 and (**ii**) −1/2 defects.

**FIG. S5:**
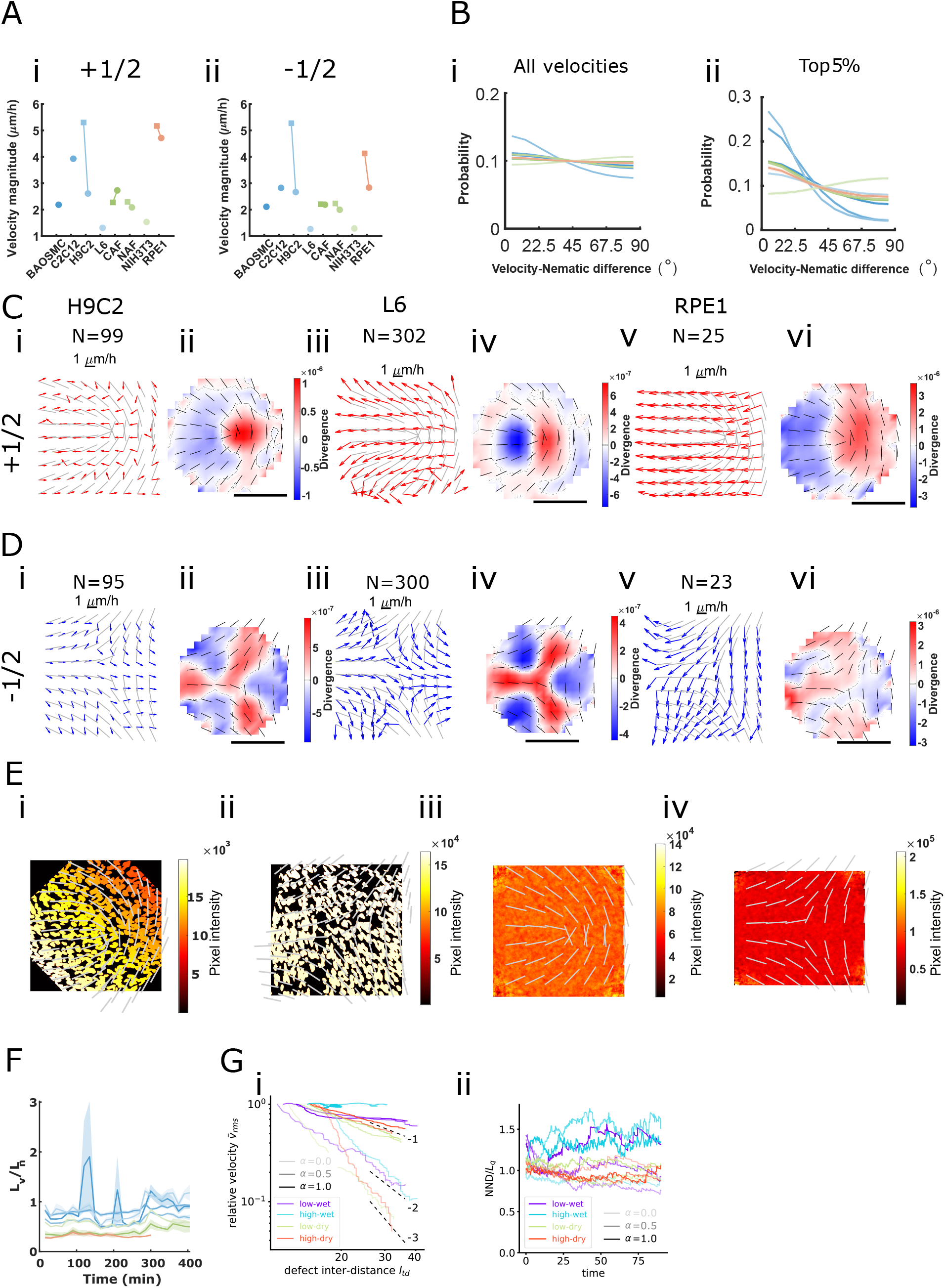
Velocity fields and fluxes around topological defects (**A**). Velocity magnitude averaged at the centre of topological defects. (**i**) +1/2 defects. (**ii**) − 1/2 defects. (**B**). Angular difference between the velocity orientation and the local nematic field orientation, measured 48 h after confluence from three independent replicates. (**i**) All spatial positions. (**ii**) Top 5% highest velocities. (**C**). Analysis of velocity fluxes around +1/2 topological defects for different cell types. (**i**-**ii**) H9C2, (**iii**-**iv**) L6, and (**v**-**vi**) RPE1 cell lines. Averaged +1/2 defect configurations showing the nematic orientation field (gray) and the averaged velocity field (red). Spatial averaging is performed using nodes of size 30 pixels (**ii**,**iv**,**vi**) Corresponding averaged divergence maps of the velocity field around +1/2 defects. (**D**). Same analysis as in (**C**) for −1/2 topological defects. (**E**). Nuclei intensity distribution around a single RPE1 topological defect, with the nematic orientation field extracted from actin-stained images. (**i**) +1/2 defects. −1/2 defects. Averaged topological defect configurations and corresponding nuclei intensity profiles are shown for (**iii**) +1/2 and (**iv**) − 1/2 defects. (**F**). Ratio of Velocity correlation length to nematic correlation length during acquisition 48h after confluence *L*_*v*_/*L*_*n*_. (**Gi**). Normalized root-mean square velocity 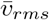 as a function of defect inter-distance *l*_*td*_ in log-log scale. (**Gii**). Ratio of nearest-neighbour defect distance (NND) to nematic correlation length (*L*_*n*_) as a function time, for the different regimes and different activities.

